# A Novel Standalone Microfluidic Device for Local Control of Oxygen Tension for Intestinal-Bacteria Interactions

**DOI:** 10.1101/2020.05.22.111096

**Authors:** Chengyao Wang, Thao Dang, Jasmine Baste, Advait Anil Joshi, Abhinav Bhushan

## Abstract

The intestinal environment is unique because it supports the intestinal epithelial cells under a normal oxygen environment and the microbiota under an anoxic environment. Due to importance of understanding the interactions between the epithelium and the microbiota, there is a strong need for developing representative and simple experimental models. Current approaches do not capture the dual-oxygen environment, require external anaerobic chambers, or are complex. Another major limitation is that in the solutions that can mimic the dual-oxygen environment, the oxygenation level of the epithelial cells is not known, raising the question whether the cells are hypoxic. We report standalone microfluidic devices that form a dual-oxygen environment without the use of an external anaerobic chamber or oxygen scavengers to coculture intestinal epithelial and bacterial cells. By changing the thickness of the device cover, the oxygen tension in the chamber could be modulated. We verified the oxygen levels using several tests: microscale oxygen sensitive sensors incorporated within the devices, hypoxic immunostaining of Caco-2 cells, and genetically encoded bacteria. Collectively, these methods monitored oxygen concentrations in devices more comprehensively than previous reports and allowed for control of oxygen tension to match the requirements of both intestinal cells and anaerobic bacteria. Our experimental model is supported by the mathematical model that considers diffusion of oxygen into the top chamber and the cellular oxygen consumption rate. This allowed us to experimentally determine the oxygen consumption rate of the epithelial cells more precisely.

## 1. Introduction

The gastrointestinal tract has a unique oxygenation profile where epithelial cells lining the mucosa and the microbiota in the lumen exist in relatively different oxygen-tension environments. Specially in the colon, the oxygen-tension varies across the lumen from anaerobic to the richly vascularized subepithelial mucosa (Albenberg et al., 2014; Karhausen et al., 2004, 2003; Sheridan et al., 1990; Zheng et al., 2015). The intestinal epithelial and bacterial cells have evolved to symbiotically exist in this metabolic environment. The commensal gut microbiota comprising facultative and anaerobic species directly (Morowitz et al., 2011; Oz, 2017) or indirectly affect important functions (Kaska et al., 2016; Melican et al., 2008; Zhao et al., 2018) such as protection against infectious pathogens, nutrient balance, metabolic homeostasis, and metabolism of drugs that can affect many disorders including Parkinson’s disease (Kessel et al., 2019; Pickard et al., 2017). Given the integral role of the epithelium and the microbiota in health, an experimental model that can mimic the environment is highly desirable (Arrieta et al., 2016; Fritz et al., 2013; Hooper et al., 2001; Nguyen et al., 2015).

There have been several attempts to recapitulate the intestinal microenvironment. Although animal models enable study of in vivo pathophysiology, the outcomes have limited translation because of the distinct differences between animal and human intestinal physiology and microbiota. As an alternative, in vitro models have utilized Caco-2, a human intestinal cell line, to represent different aspects of the intestinal physiology such as barrier function, drug transport, and interactions with bacteria. The culture methods have evolved from the traditional Transwell inserts which suffered because they could not properly separate and monitor the culture chambers into different oxygen-tension environments (Dutta and Clevers, 2017; Fatehullah et al., 2016; Sadaghian Sadabad et al., 2015). For instance, Ulluwishewa *et al*. modified the apical chamber of Transwell inserts to make it anaerobic and to co-culture Caco-2 cells with live *Faecalibacterium prausnitzii* (Ulluwishewa et al., 2015), however, the oxygen levels in the Transwell chambers were not monitored. Kim *et al*. developed a self-sustaining oxygen gradient platform to study the interaction of the crypt base and obligate anaerobes by incorporating a polycarbonate plug in a Transwell insert (Kim et al., 2019). Other strategies use 3D culture models like sacrificial micromolding (Samy et al., 2019) and microfluidic tissue culture arrays (Beaurivage et al., 2019) to mimic the intestinal-bacterial environment. The basic limitation of these approaches is that there is a lack perfusion of nutrients to the cells, which has been shown to be important for cell function and maintenance of tight junctions (Kim et al., 2016).

The use of microfluidics in the recent intestinal organ-on-chip models (Kim et al., 2016, 2012; Pearce et al., 2018; Shah et al., 2016) has enabled construction of chambers with different oxygen concentrations to co-culture bacteria and Caco-2 cells in anaerobic and aerobic conditions (Jalili-Firoozinezhad et al., 2019). S. Jalili-Firoozinezhad *et. al* placed the device into an anerobic chamber while perfusing oxygenated media to establish two oxygen tensions within the 2-layer device to coculture Caco-2 cells with microbial samples as well as integrated microscale oxygen sensors into the device for in situ oxygen measurements (Jalili-Firoozinezhad et al., 2019). Another approach achieved control of oxygen (Shin et al., 2019), however, maintenance of the oxygen gradient required continuous perfusion of the anoxic media which limited the cell-bacteria interactions as shown by others (Jalili-Firoozinezhad et al., 2019). Using a different approach, Gao et al. used oxygen scavengers on a non-permeable material to control two different oxygen tensions, however, because of a lack of membrane, the area of interaction between the cells in the normoxic and hypoxic chambers was hindered (Gao et al., 2019).

Although these strategies were able to achieve a dual oxygen-tension environment, several critical questions remain unanswered. Since the interface between the bacteria and the epithelial cells is just a few microns thick, separated by only the mucus layer, the hypoxia state of the intestinal cells was not determined in these studies. Without knowing the oxygenation-state of the epithelial cells, it was impossible to determine whether the experimental model actually mimics the in vivo intestinal environment. This is important because we know that if hypoxia in the intestinal cells is not controlled, cellular function can become impaired(Glover and Colgan, 2011; Lei et al., 2014; Melican et al., 2008; Sturrock et al., 2018; Xu et al., 1999; Zeitouni et al., 2016). In addition, these devices are quite complex to fabricate and operate, or use synthetic oxygen scavengers which are cytotoxic and take considerable time to establish the oxygen gradient. Some approaches still required placing the device in an anaerobic gas chamber to achieve the oxygen gradient (Jalili-Firoozinezhad et al., 2019), whereas others rely on a continuous flow of deoxygenated media to maintain the oxygen gradient which limited the interactions between bacteria and intestine (Jalili-Firoozinezhad et al., 2019).

To address these gaps, we have developed a novel microfluidic device which leveraged the permeability of PDMS to create dual-oxygen environment in our intestine-on-a-chip device; in other words, the intestinal epithelial cells were in normoxic conditions whereas the bacteria were under anoxic conditions. We verified the oxygen status by three complementary approaches – (1) oxygen levels in the device were measured by incorporating oxygen sensors in situ, (2) oxygen levels of the intestinal cells were determined by pimonidazole, and (3) oxygen levels of the bacteria were determined by measuring growth of a genetically modified a facultative *E. coli* strain. We then mathematically modeled the oxygen transport across the cellular layer and experimentally determined the oxygen consumption rate of the intestinal cells during culture with bacteria, which to our knowledge has never been reported before.

## 2. Methods and Materials

### 2.1 Microfluidic Device Fabrication

The basic device builds upon our previous work in which a microfluidic device with two chambers separated by a porous membrane was developed (Hegde et al., 2014; Wang et al., 2018). Briefly, SYLGARD 184 Silicone Elastomer Kit containing silicone elastomer base and curing agent (Dow Chemical) was used to fabricate the polydimethylsiloxane (PDMS) devices. All device chambers were created using soft lithography using a lithographically patterned SU-8 wafer mold. The cell culture channel in the device was 1 mm wide, 1 cm long, and 100 μm tall. To fabricate the devices, the base and curing agent were mixed at ratio of 9:1 and poured into the mold. The molds were placed in a vacuum chamber (Bel-Art Products) overnight to remove any air bubbles and then placed in a 65°C oven overnight for curing. A biopsy punch (World Precision Instruments) was used to cut 0.75 mm holes to provide access for tubing (Cole-Parmer) that has inner diameter of 0.25 mm and outer diameter of 0.75 mm. For the 1-chamber device, the device was bonded to a clean glass slide using a plasma cleaner (Harrick Plasma) (**Figure 1A**). The resulted 1-chamber device includes a chamber with volume of 1 μL. To fabricate the 2-chamber device, first a 250 μm thick silicone sheet (Rogers Corporation) and a porous polyester membrane (AR Brown-US) were cut using a laser cutter (Epilog Laser, CO) and cleaned with 70% isopropyl alcohol (Fisher Scientific). The porous polyester membrane had pore size of 0.4 μm, pore density of 4×10^6^ pores/cm^2^ and thickness of 10 μm (AR-Brown). The silicone sheet was bonded to a clean glass slide using the plasma cleaner. A porous membrane was attached to the PDMS device (single chamber above) using SYLGARD 184 Silicone Elastomer, cured overnight in 65°C oven, and bonded to the siloxane sheet on the glass slide using the plasma cleaner (**Figure 1B**). The resulted devices include a top chamber with volume of approximately 1 μL and a bottom chamber with volume of approximately 2.5 μL. For both devices, media reservoirs were created by cutting out 8 mm holes from a separate PDMS layer and bonded to the top of the devices using plasma cleaner. The dead volume of tubing for connecting flow was 50 μL. The total surface for cell seeding in the both devices was 0.1 cm^2^. The device chambers were cleaned and sterilized with 70% isopropyl; the surface of device was sterilized under UV light for 15 minutes for each side before use. The distance that devices were put from the UV light source is 50-100 cm.

**Figure 1:**
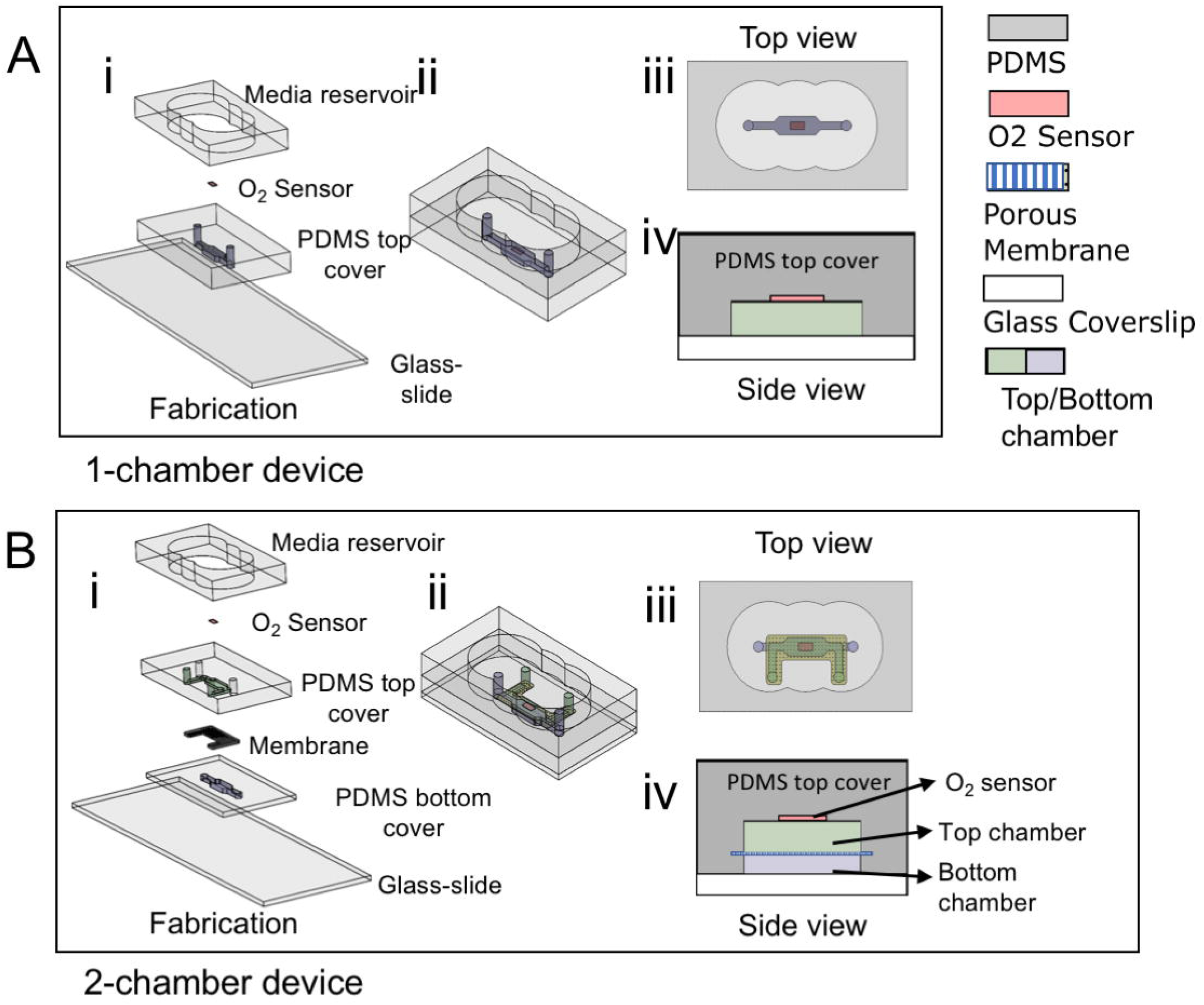
Schematic of fabrication process for 1-chamber and 2-chamber devices. (A) 1-chamber device: (i) the 3D view of fabrication process; (ii) 3D view of fabricated device; (iii) the top view and (iv) the side view of fabricated devices. (B) 2-chamber device with porous membrane: (i) the 3D view of fabrication process; (ii) 3D view of fabricated device; (iii) the top view and (iv) the side view of fabricated devices.

Additional PDMS was added to the top of the devices to create a ‘top cover’. Devices with two different top cover thicknesses were made – i) standard device in which the thickness of the top cover was 2.5 mm and ii) thick device in which the top cover was 5 mm thick.

### 2.2 Oxygen Sensor Fabrication

Oxygen sensors were made using Pt(II)*meso*-tetrakis(pentafluorophenyl)porphine (PtTFPP) (Frontier Scientific) using a method described previously (Thomas et al., 2009). Briefly, 11.7 mg PtTFPP was dissolved in 2 mL toluene and then mixed with 10 g PDMS (base and curing agent mixed at 9:1), spun on a glass slide to form an approximately 150 μm thick layer and then cured in 65°C overnight. The film was cut out and placed on the patterned wafer mold to overlap the device chamber. Previous work confirmed that a PDMS film with PtTFPP particles was suitable for cell culture and did not affect the cellular behavior (Jenkins et al., 2015; Thomas et al., 2009).

### 2.3 Mathematical Modeling of Oxygen Transport

There are two sources that bring oxygen into the cell culture chamber. One, oxygen from the environment that will diffuse into the device through the top cover. The second source is from the oxygen in the perfused media that can diffuse through the membrane and through the cell monolayer (the unconsumed portion of oxygen).

#### a) Diffusion of oxygen from the environment

We modeled the transport of oxygen through the device top cover (**Figure 2**). Using Fick’s Second law of diffusion which relates concentration of oxygen with time and thickness across the device top cover (*Eq. 1*), we evaluated the concentration of oxygen diffusing through different thicknesses of the top cover under steady-state (*Eq. 2*) (Bird et al., 2007).

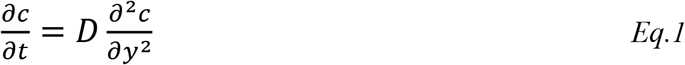

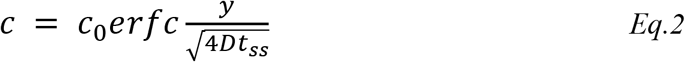

where, *c* is the concentration of oxygen in device and *c*_0_ is the concentration of oxygen in the atmosphere outside the device and was taken to be constant at 21%, and *D* is the oxygen diffusion coefficient in PDMS. The distance from PDMS top surface to chamber is *y* and *t_ss_* is the time it takes for oxygen concentration to reach steady-state in PDMS (Thomas et al., 2011). The parameters used in the mathematical model are listed in **Table. 1**.To experimentally validate the mathematical model, cells and bacteria will be cultured in the 1-chamber device with different top cover thicknesses (**Figure 3A-B**). In this work, we defined the environment hypoxic when the oxygen concentration was less than 1% and normoxic when it was equal to the oxygen concentration in incubator, which was 18.5% (Newby et al., 2005; Place et al., 2017).

**Figure 2:**
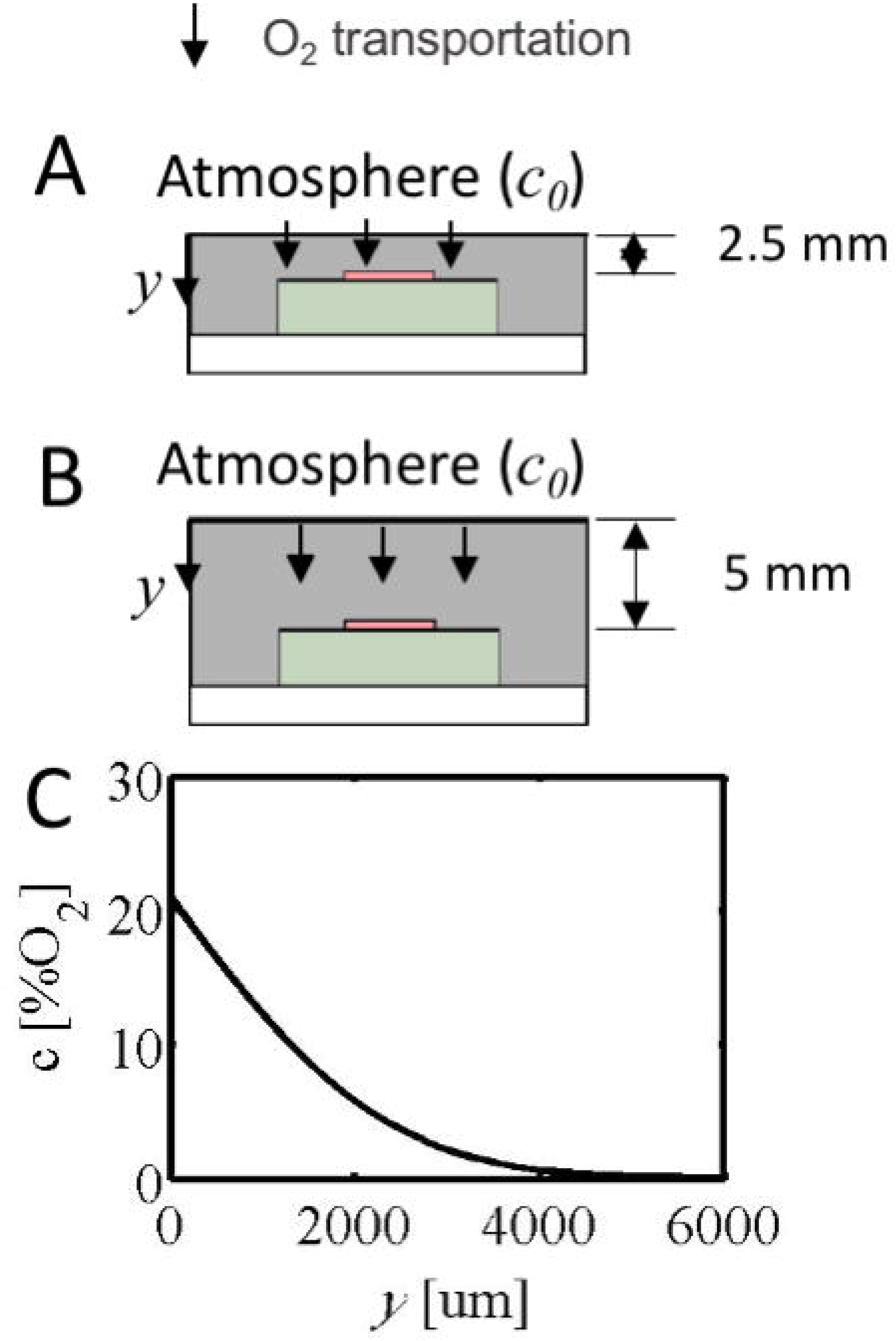
(A-B) Side view of 2.5mm thick (A) and 5mm thick (B) top cover devices. (C) 1-D Fick’s law simulation model of oxygen transport into the device through the top cover.

**Figure 3:**
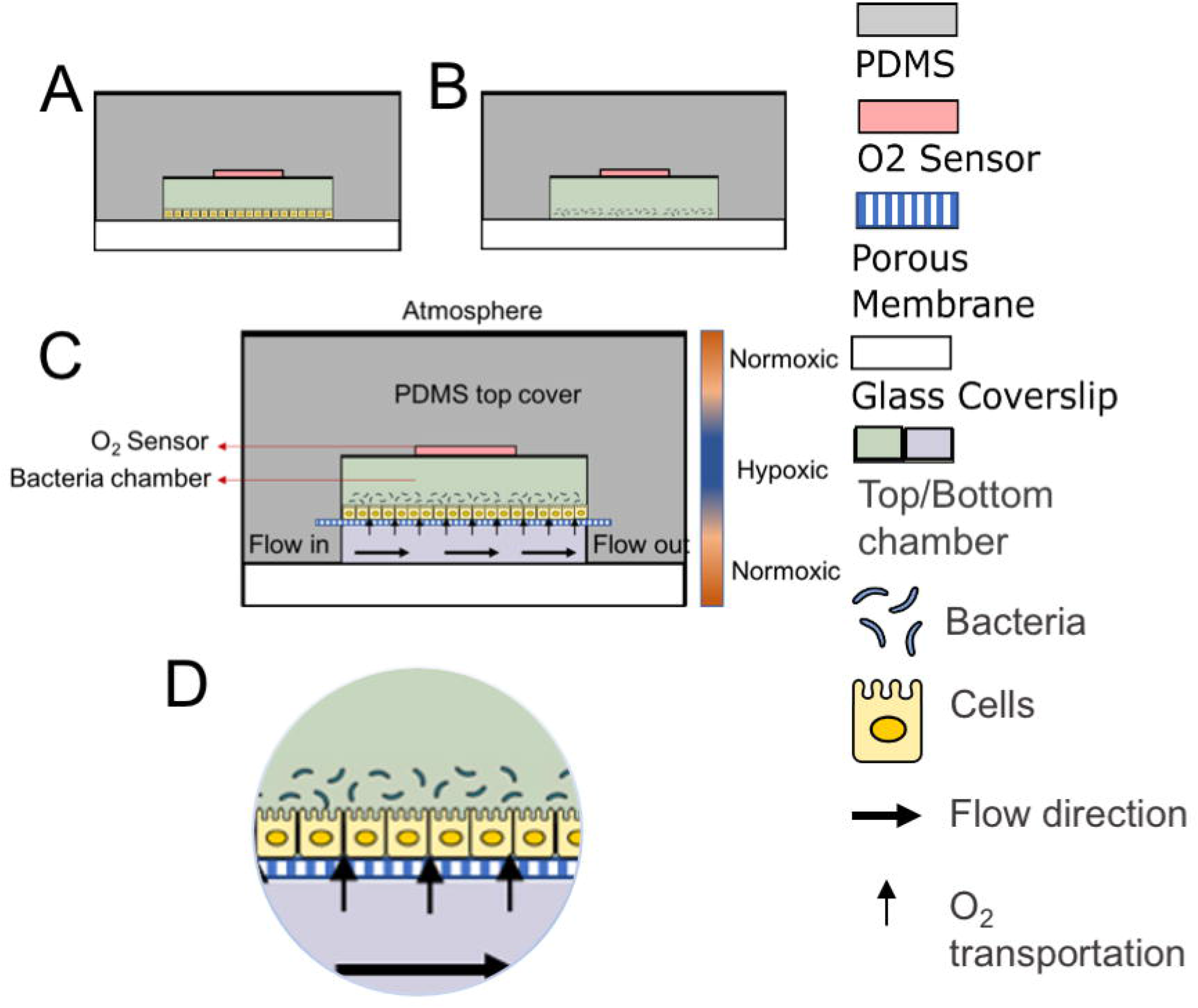
(A-B) Side view of Caco-2 cells (A) and bacteria (B) seeded in 1-chamber device. (C) Side view of Caco-2 cells and bacteria co-cultured in top chamber of a 2-chamber device, with media in the top chamber static and that in the bottom chamber perfused. (D) Detailed side view of co-cultured bacteria and cells on porous membrane, cell culture medium flow direction and oxygen transport through the membrane. (E) F-actin (Red) and Hoechst (blue) stained Caco-2 cells imaged on day 3. Images were stitched together to show a fully confluent monolayer of intestinal epithelial cells along the entire device. Scale bar: 200 μm.

**Table 1.** Parameters of mathematical modeling of Eq. 2

| Parameter | Value |
| --- | --- |
| $D$ (diffusion coefficient of O<sub>2</sub> in PDMS) | $3.25 \times 10^{-9}$ m<sup>2</sup>/s (Markov et al., 2014) |
| $t_{ss}$ (steady-state time of oxygen transportation in PDMS) | 510 s (Thomas et al., 2011) |
| $c_0$ (% O<sub>2</sub>) | 21 |

#### b) Transport of oxygen from the perfused chamber to the top

Since the one-chamber devices do not have a membrane, this analysis was carried out only for the 2-chamber devices. In this case, to estimate the oxygen concentration in the upper chamber, in addition to the transport of oxygen into the top cover, we accounted for the (1) diffusion of oxygen from the media in the bottom chamber through the porous membrane and (2) oxygen consumed by the cells (**Figure 3C-D**). In this model, we considered a confluent monolayer of Caco-2 cells in the top chamber; media was perfused in the bottom chamber. The Caco-2 monolayer was monitored daily under a microscope (Nikon ECLIPSE Ti) until confluent (**Figure S1**). The cell count in the confluent cell layer was quantified by staining the nuclei with a nuclear stain and counting the nuclei using ImageJ (NIH) – for our devices, a confluent layer comprised 0.0215 ± 0.003 × 10^6^ Caco-2 cells, which means the density was 0.215±0.03 × 10^6^ cells/cm^2^. Oxygen consumption by the cells lowered the concentration of oxygen in the top chamber. Our objective was to keep the intestinal epithelial cells normoxic while keeping the bacteria that sit above the cells anoxic. We used the model to determine whether the consumption of oxygen by the cells would make the top chamber environment for the bacteria anoxic and to determine the rate-limiting flow rate at which this would happen.

To determine the transport of oxygen from the media perfused in the bottom chamber through the membrane, we first calculated the Péclet Number, which came out to be 0.005 (≪ 1), indicating that diffusion was the primary mode of transport for oxygen from the bottom chamber to top chamber. Using Darcy’s law, we determined that the transport of oxygen through the membrane would take place at approximately 1/100 of the perfusion flow rate in the bottom chamber (Chung et al., 2018, 2014; Loskill et al., 2017).

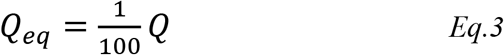

where, *Q_eq_* is the equivalent flow through the membrane and *Q* is the flow rate of perfusion in the bottom chamber.

We also calculated the oxygen consumed by the cells. The only oxygen available to the cells was the oxygen transported through the membrane. We defined *q* as the oxygen consumption rate of a single Caco-2 cell(Decleer et al., 2018; Grauso et al., 2019; Zeitouni et al., 2015) (which was calculated in our model), and *N* as the total number of Caco-2 cells in the device. We can estimate the flow rate of a fluid, *Q_o2_*, that contains oxygen at a concentration of *c_m_* which would balance the oxygen requirements of the cells (*Eq. 4*). We estimated *N* to average 0.0215 × 10^6^ cells and *c_m_* was taken as 2.1 × 10^-3^ mol/L (Zeitouni et al., 2015).

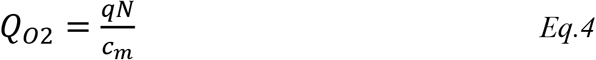

For the case when the oxygen transported through the membrane will be equal to the oxygen consumed by the cells, we calculated the flow rate, *Q*, that would be required to provide sufficient oxygen to the cells (**Table 2**) and was experimentally validated as described below.

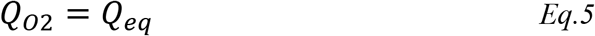

**Table 2.** Flow rate calculated from Eq. 3 using oxygen consumption rate of Caco-2 cells (Zeitouni et al., 2015),(Decleer et al., 2018),(Grauso et al., 2019).

| Consumption rate $q$ (mol/s · cell) | | Calculated flow rate $Q$ (μL/h) | |
| --- | --- | --- | --- |
| from | to | from | to |
| $0.34 \times 10^{-17}$ | $2.08 \times 10^{-17}$ | 0.30 | 1.82 |
| $0.04 \times 10^{-17}$ | $0.18 \times 10^{-17}$ | 0.04 | 0.16 |
| $3.5 \times 10^{-17}$ | | 3.06 | |

### 2.4 Epithelial Cell culture

Human intestinal epithelial cells Caco-2 (ATCC^®^ HTB-37™) were cultured in a 75 cm^2^ cell culture flask (Nest Biotechnology) with complete 1X Eagle’s Modified Eagle Medium (EMEM) (VWR) which comprised of 4.5 g/l glucose, 25 mM HEPES, 10% Fetal Bovine Serum (FBS) (Gibco), and 1% pen-strep (Gibco). The cells were cultured until they reached 70% confluency prior to passaging. Cells were incubated at 37°C and 5% CO_2_ in a humidified incubator and media was changed every other day. Before seeding into the device, cells were counted and observed using Trypan Blue (Sigma Aldrich) under microscope (Zeiss Axiovert 40C, Thornwood, NY). The sterilized device was coated first with 50 μg/mL Bovine Plasma Fibronectin (Gibco) and then with 100 μg/mL Type I collagen (BD); each for 30 minutes and at 37°C. Caco-2 cells were seeded at density of 2×10^4^ cells/cm^2^. After allowing 3-hours for the cells to attach, fresh cell culture media was perfused for up to four days.

### 2.5 Bacteria Culture

As a facultative bacteria, *E.coli* Nissle 1917 (ECN) (Ardeypharm GmbH, Germany) can survive in both anoxic and normoxic environments but exhibits a slower growth rate in anoxic environments (Sonnenborn and Schulze, 2009). *Bifidobacterium adolescentis* Reuter (*B. adolescentis*) (ATCC^®^ 15703) is an anaerobic bacterium due to which its growth is hampered in an aerobic environment as compared to that in an anaerobic environment. The growth of these two bacteria can be used to determine the presence of anaerobic/aerobic environments.

ECN was cultured in a conical flask with 50 mL of Luria-Bertani (LB) broth overnight. The LB broth for ECN consisted of 1% (W/V) Tryptone (Fisher BioReagents), 0.5% (W/V) yeast extract (Fisher BioReagents), and 1% (W/V) sodium chloride (Columbus Chemical Industries). The culture was grown in an aerobic environment at incubation temperature of 37°C and with shaking speed of 200 rpm.

A separate culture of ECN was labeled with Green Fluorescent Protein (GFP) using methods described before (Fang et al., 2018; Lizier et al., 2010; Schultz et al., 2005). The labeled bacteria were selected using 0.001% (v/v) erythromycin (Fisher BioReagents). The GFP labeled ECN showed increasing fluorescence intensity while they were growing.

*B. adolescentis* was cultured in a 50 mL conical flask with 3.8% (W/V) Difco modified reinforced clostridial medium (BD) for 48-hours in an anaerobic jar. The anoxic environment was made by using the GasPak EZ™ Anaerobe Container System with Indicator (BD) as instructed then incubated at 37°C under a shaking speed of 200 rpm. The GasPak EZ™ Anaerobe Container System can create less than 1.0% oxygen, and greater than or equal to 13% carbon dioxide.

For both strains of bacteria, the standard curve of the bacteria number and optical density (OD) was plotted first (**Figure S2**) and then the growth was quantified using the optical density (OD) according to the standard curve we made.

### 2.6 Culture conditions

All experiments were performed in either a conventional aerobic humidified incubator with 5% CO_2_ or an anaerobic environment which were created by using BD GasPak EZ anaerobe container system. Based on the determination of the hypoxia and normoxia mentioned above, we defined the first one as normal culture condition and second one as hypoxic culture condition. The device with top cover thickness 2.5 mm in the environment with normal oxygen (Std-O_2+_), was used as the control. The hypoxic environments include: device with top cover thickness 2.5 mm in anaerobic chamber (anoxic) (Std-O_2-_), device with top cover thickness 2.5 mm with oxygen scavenger pyranose-2-oxidase (P2O)(Jung et al., 2018) (anoxic) (Std-P2O), device with top cover thickness 5 mm in environment with normal oxygen (Thick-O_2+_), device with top cover thickness 5 mm in anaerobic chamber (anoxic) (Thick-O_2-_), device with top cover thickness 5 mm with oxygen scavenger (Thick-P2O). The experimental groups are listed in **Table 3**.

**Table 3.** Experimental groups of devices, different culture conditions and bacteria growth.

| Experimental Groups | Experimental condition |  | Hypoxic condition or not | *B. adolescentis* growth | ECN growth |
| --- | --- | --- | --- | --- | --- |
|  | Top cover thickness | Culture condition |  |  |  |
| Std-O<sub>2</sub><sup>+</sup> | 2.5mm | Environment with normal oxygen (normoxic) | No | down | up |
| Std-O<sub>2</sub><sup>-</sup> | 2.5mm | In anaerobic chamber (anoxia) | Yes | up | down |
| Std-P2O | 2.5mm | With oxygen scavenger (anoxia) | Unknown | down | down |
| Thick-O<sub>2</sub><sup>+</sup> | 5mm | Environment with normal oxygen (normoxic) | Yes | up | down |
| Thick-O<sub>2</sub><sup>-</sup> | 5mm | In anaerobic chamber (anoxia) | Yes | up | down |
| Thick-P2O | 5mm | With oxygen scavenger (anoxia) | Unknown | down | down |

### 2.7 Oxygen Sensor Calibration

Different oxygen tensions were applied to the oxygen sensor to calibrate the films. The oxygen sensors were imaged using fluorescent microscope (Nikon ECLIPSE Ti, Tokyo, Japan), and the intensity was calculated using ImageJ (NIH). The calibration curve is shown in **Figure S3**.

The standard 1-chamber devices with oxygen sensors embedded were equilibrated in room condition, standard cell culture incubator and anaerobic jar for 24 hours; oxygen concentration in each condition was considered to be 21%, 18.5% and 0% respectively (Newby et al., 2005; Place et al., 2017). The emission intensity of oxygen sensor in different devices was recorded every second till the fluorescent intensity became stable by the fluorescent microscope. All images were analyzed using ImageJ software. The fluorescent intensity was determined through the Stern-Volmer equation (Amao et al., 2001; Yeh et al., 2006).

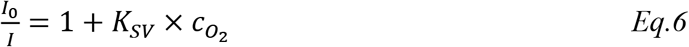

where, *I*_0_ is the intensity of in the absence of oxygen (in anaerobic jar) and *I* is the intensity of sensors in the presence of oxygen. *C_o_2__* is the oxygen concentration in different conditions. *K_SV_* is the Stern-Volmer constant. The intensities were obtained and plotted (**Figure S3**).

### 2.8 Hypoxia test

To test the growth of the aerobe and anaerobe in the devices, we cultured each species in the 1-chamber devices. Before seeding, the optical density (OD) was measured using the Spectra Max M2 (Molecular Devices, San Jose, CA) at an absorbance wavelength of 600 nm. The result was then compared to the characteristic standard curve (**Figure S2**) of each bacteria species to quantify the density of seeding. Bacteria was diluted and directly seeded into sterile device at 0.1 x 10^6^ CFU/mL. The diluted bacteria suspensions were then serially diluted as 1/1, 1/10, 1/25 and 1/50 before spread on agar plate as control. ECN and GFP-ECN were similarly cultured in 1-chamber devices for 24 hours and *B. adolescentis* for 48 hours. After seeding, the inlets and outlets of chamber were sealed with tubing. Both standard and thick devices were placed in normoxic and anoxic environments. After 24 hours and 48 hours, ECN and *B. adolescentis* were imaged (**Figure 4A** and **4C**) using a brightfield microscope (Nikon ECLIPSE Ti, Tokyo, Japan). GFP-ECN in device was imaged (**Figure 4E**) using a laser scanning confocal microscope (Carl Zeiss Microscopy, Thornwood, NY). ECN were eluted out of the devices and spread on LB agar plate (Fisher BioReagents) for 24 hours; and *B. adolescentis* were eluted out and spread on Reinforced Clostridial agar plate (OXOID) for 48 hours. Both bacteria suspensions eluted out were diluted as 1/50 before spread on the agar plates. The ECN plates were placed in a standard bacteria incubator, while the *B. adolescentis* plates were placed in the GasPak EZ anaerobe container system. The bacteria colonies on the plates were imaged and counted using ImageJ. The relative change of bacteria growth in device to initial bacteria seeding density was determined (**Figure 4B** and **4D**). For GFP-ECN, we obtained the mean intensity of fluorescence by using LSM5 Pascal program (Zeiss) and plotted in **Figure 4F**. For each test, the replicas n ≥ 5.

**Figure 4:**
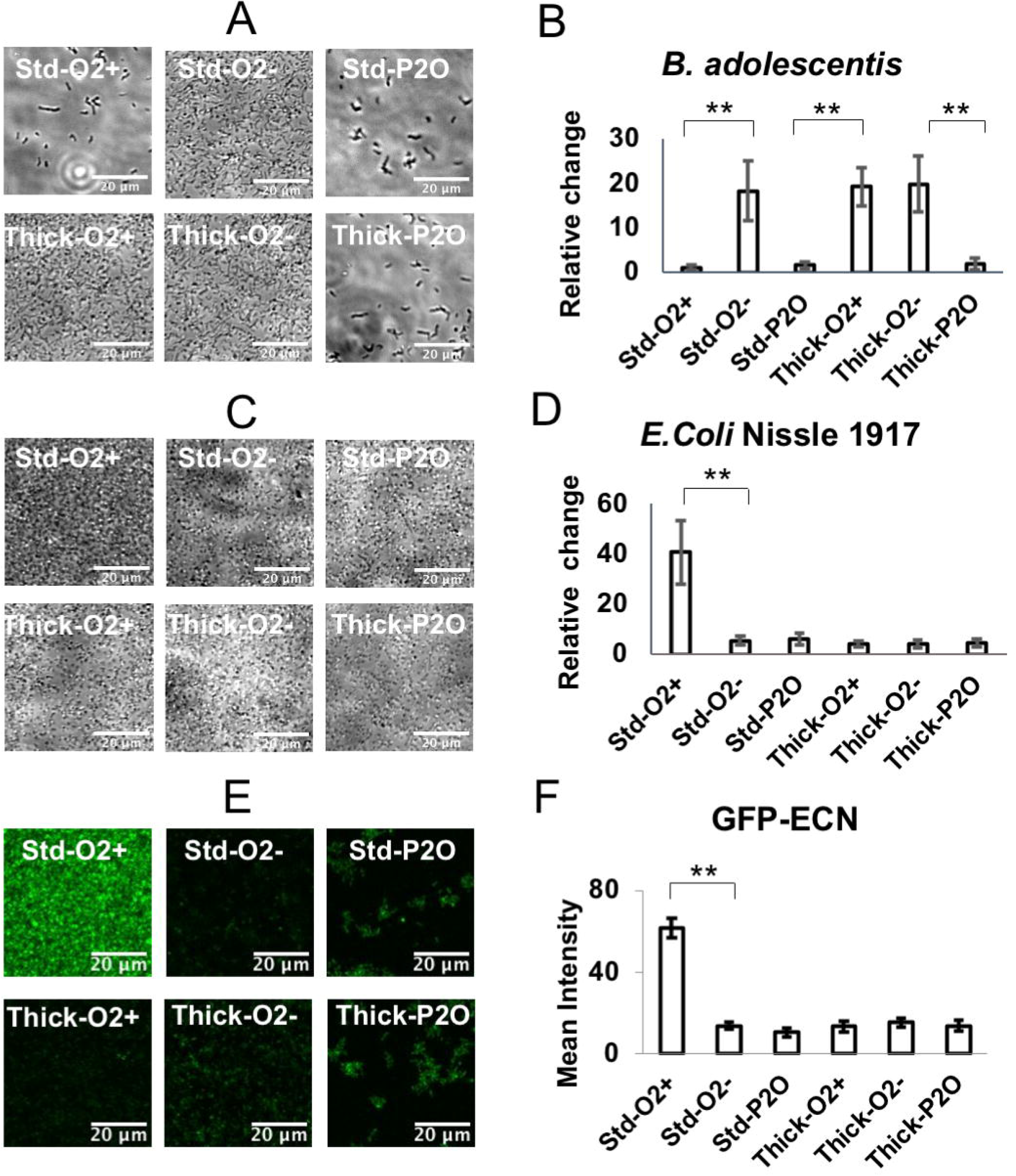
(A) The growth of anaerobic bacteria – *B. adolescentis* in standard (2.5mm) and thick (5mm) PDMS top covered 1-chamber device for 48 hours in normoxic or anoxic culture condition. Scale bar is 20 μm. (B) The growth quantification of *B. adolescentis* in (A). (C) *E.coli* Nissle 1917 in standard and thick PDMS top covered 1-chamber device for 24 hours in normoxic or anoxic culture condition. Scale bar is 20 μm. (D) The growth quantification of *E.coli* Nissle 1917 in (C). (E) Fluorescent images of GFP labeled *E.coli* Nissle 1917 (GFP-ECN) growth in standard and thick PDMS top covered 1-chamber device for 24 hours in in normoxic or anoxic culture condition. Scale bar is 100 μm. (F) Mean fluorescence intensity quantification of GFP-ECN growth in (E). *: *p*-value < 0.05; **: *p*-value < 0.01.

To validate hypoxia in Caco-2 cells, we cultured the cells in different tissue culture plates and placed the plates in different environments: (a) standard tissue culture incubator and (b) anaerobic jar. After 24 hours, the cells were treated with 200 μM pimonidazole HCl (Hypoxyprobe) for 2 hours to determine whether the cells were hypoxic(Bao et al., 2012). Pimonidazole HCl reduces in hypoxic cells to form stable covalent adducts with thiol groups that can be detected under a fluorescent microscope (Aguilera and Brekken, 2014; Raleigh et al., 1998; Raleigh and Koch, 1990). After pimonidazole treatment, the cells were fixed and stained with FITC-Mab (Hypoxyprobe) and FITC HRP MAb (Hypoxyprobe) and Hoechst (Thermo Fisher). The result of staining was then imaged using Nikon fluorescent microscope (**Figure S4**). For testing hypoxia in cells in the microfluidic devices, Caco-2 cells in the 1-chamber device with standard or thick PDMS top layers were exposed to hypoxic conditions for 0, 2, 4, 8, and 24 hours before staining using pimonidazole as described (**Figure 5**, **Figure S5**). To quantify the hypoxia expression, fluorescence intensity and cell number in each image was obtained through Image J. To normalize the expression, Caco-2 cells cultured in device without hypoxic treatment, which is 0 hour of each group, was used as the control group.

**Figure 5.**
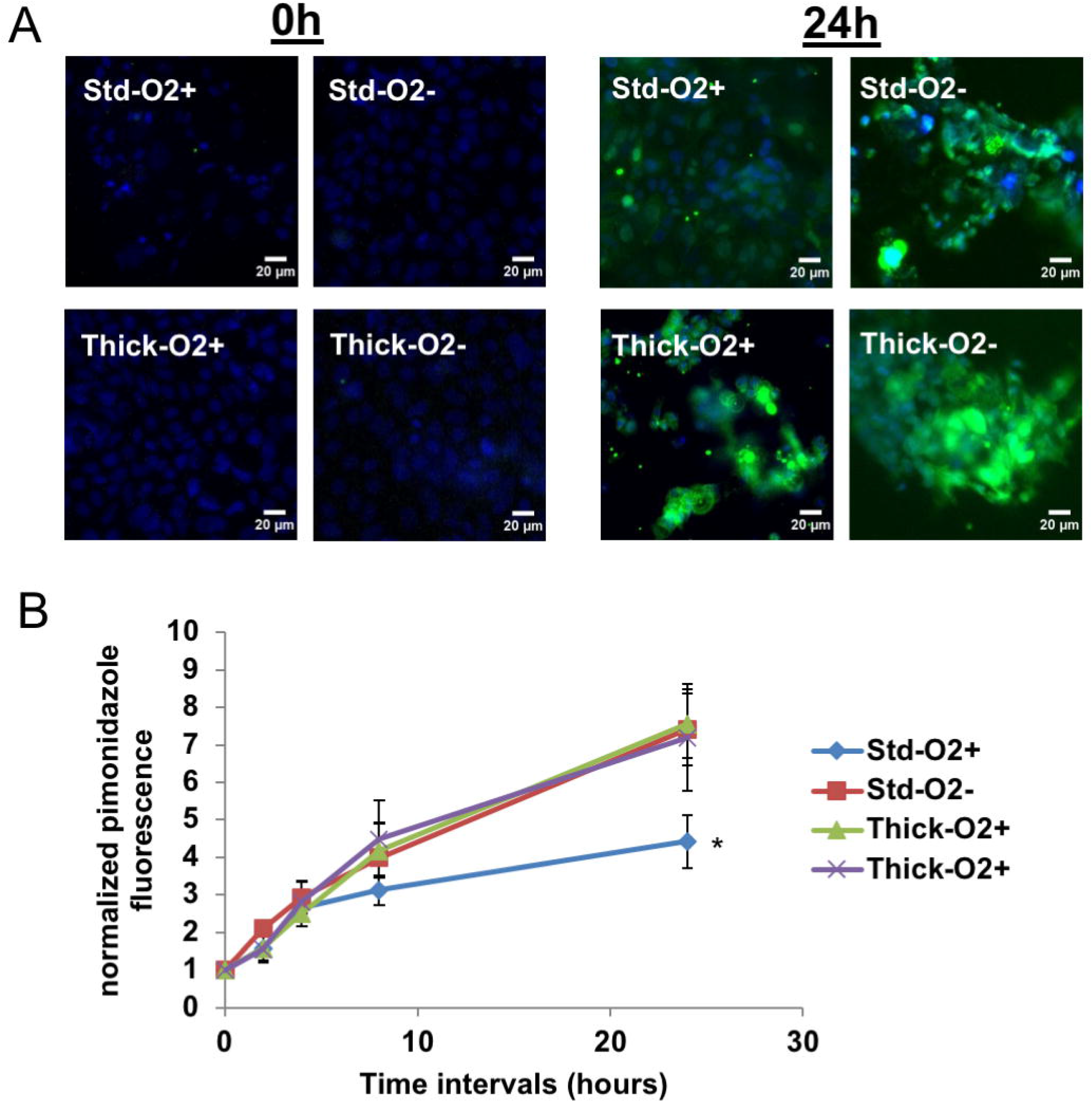
(A) Hypoxia immunostained images of normoxic and hypoxic Caco-2 cells in devices with standard and thick top cover for different treatment intervals: 0 and 24 hours, using pimonidazole staining kit. (B) The quantification of hypoxia in the cells in devices with standard and thick top cover for different treatment intervals: 0, 2, 4, 8 and 24 hours. Statistical significance was tested using one-way ANOVA. Green: hypoxia; Blue: nuclei. Scale bar is 20 μm. *: *p*-value < 0.05; **: *p*-value < 0.01.

To study the intestine-bacteria interaction, we cocultured bacteria and Caco-2 cells in the 2-chamber device with thick top cover and in normal incubation conditions (Std-O_2+_). The thick device was used to create an anoxic environment in top chamber as per the mathematical model described above. Caco-2 cells were seeded into the top chamber. Oxygen was monitored using three different methods – (1) in the top chamber, using the in-situ oxygen sensors, (2) pimonidazole for the Caco-2 cells, and (3) GFP for the bacteria. Caco-2 cells attached and became confluent after approximately four days of perfusion. The confluency of Caco-2 monolayer was monitored and the F-actin was stained using phalloidin (Invitrogen) every day before cells were confluent. 24 hours before co-culture, the Caco-2 cells were perfused with antibiotic free EMEM medium. Before bacterial seeding, the optical density (OD) was measured as indicated above and the result was then compared to the standard curve (**Figure S2**) of each bacteria species to quantify the multiplicity of infection (MOI). The MOI we used in this study is the ratio of bacteria: Caco-2 cells = 10:1. Then *B. adolescentis* and ECN were mixed in antibiotic free EMEM media and seeded over the confluent Caco-2 cells. The top chamber was sealed while the bottom chamber was then perfused with antibiotic free EMEM media after bacterial seeding. The perfusion rate was varied from device 5, 7.2 to 30 μL/h based on the oxygen diffusion simulation to investigate different oxygen gradients. The devices underwent regular incubation for 24 hours. The oxygen concentration in top chamber was quantified using the oxygen sensors in device (**Figure 7A-C**, **Table S1-S3**). The hypoxia of Caco-2 cells was determined using Pimonidazole and the *B. adolescentis* growth was then quantified by spreading bacteria on agar plate. In each flow rate, the group that cells were not cocultured with bacteria was set as hypoxia expression control. To normalize the anaerobe growth in cocultured device, we used the initial bacteria seeding density as control and analyze the growth based on that. For each test, the replicas n ≥ 6.

### 2.9 Immunofluorescence imaging

#### Pimonidazole

After the cells were treated with 200 μM pimonidazole HCl, cells were fixed with 3.7% paraformaldehyde (PFA) for 10 min. After fixation, the cells were permeabilized with 0.1% (v/v) triton X100 (Sigma Aldrich) in phosphate-buffered saline (PBS) (Gibco) solution for 10 minutes at room temperature. The cells were then washed with PBS three times. A 1/100 (v/v) dilution was made with the primary antibody, FITC-MAb (Hypoxyprobe) and 1% Bovine Serum Albumin with PBS (BSA-PBS). The solution was added into the devices and left overnight at 4°C. After 24 hours, PBS was used to wash out the antibody three times. The same dilution was made with the secondary antibody, anti FITC HRP MAb (Hypoxyprobe). The solution was added for 1 hour at room temperature. After the secondary antibody was incubated and washed out, 1/1000 (v/v) dilution of Hoechst (Thermo Fisher) was added for 10 minutes at room temperature to stain the nuclei. The cells were then washed with PBS and imaged with the Nikon fluorescent microscope and the intensity calculated by ImageJ.

#### F-actin and ZO-1

Following fixation, all cells were washed with 1X PBS and permeabilized with 0.25% (v/v) Triton X100-PBS for 10 minutes at RT. After washing, the cells were blocked with 3% (v/v) BSA-PBS at RT for 30 minutes. To stain F-actin, Phalloidin was added into device and incubated at room temperature for 1 hour. To stain the tight junctions, the cells were incubated with ZO-1 antibody at 4°C overnight, after which the cells were washed with PBS and the secondary antibody was added into device and incubated at room temperature for 1 hour. The nuclei were stained as mentioned above.

The cells were imaged using a fluorescent microscope. The fluorescence intensity and nuclei number were recorded. The cell number was counted by nuclei staining and the fluorescence intensity per nuclei was quantified by ImageJ (NIH). This data was normalized using the data from the devices without treatment.

### 2.10 Data analysis

Unless otherwise indicated, data are expressed as means ± SEM. The replicas of each experimental test are n ≥ 3. Differences between experimental groups were compared using oneway ANOVA and t-test. To avoid of statistically falling into type I and type II errors increasing during one-way ANOVA and multiple t-test, we used Bonferroni correction and Holm method. Statistical analyses were done with Excel. Differences with a *p* value less than Bonferroni corrected α were considered statistically significant.

## 3. Results and discussions

### 3.1 Mathematical simulation result of oxygen diffusion in device

The MATLAB simulation result showed that when the PDMS top cover was less than 2.5 mm, the oxygen concentration in the top chamber was > 5% (**Figure 2**). In contrast, when the top cover was >5 mm thick, the oxygen concentration in the top chamber was less than 0.1% or hypoxic. Based on these results, we designed our device with a 5 mm top cover to limit the diffusion of atmospheric oxygen into the top chamber.

To model the diffusion of oxygen in the 2-chamber device (**Figure 3**), the oxygen consumption rate of Caco-2 cells in top chamber and the amount of normal oxygen cell culture medium diffused from bottom chamber to top chamber were compared. We then calculated the flow rate, *Q*, that would be required to provide sufficient oxygen to maintain the cells as normoxic. As shown in **Table 2**, there is no good agreement in the values reported for oxygen consumption rate by these cells, which is likely due to the different culture conditions. Based on these results and preliminary experiments, we selected 5 μL/h, 7.2 μL/h, and 30 μL/h as flow rates to test.

### 3.2 Control of oxygen levels in 1-chamber device

Since *B. adolescentis* is an anaerobe, it grows well in anoxia as compared to the growth under normoxic conditions. In our experiments, *B. adolescentis* had higher growth in the Std-O_2-_, Thick-O_2+_, and Thick-O_2-_ conditions (**Figure 4A**) compared to that in Std-O_2+_, Std-P2O, and Thick-P2O conditions (**Figure 4B, Table S1**). The relative change of *B. adolescentis* increase in the Std-O_2-_, Thick-O_2+_, and Thick-O_2-_ conditions, was significantly higher compared to the Std-O_2+_, Std-P2O, and Thick-P2O devices. Since the growth of *B. adolescentis* was higher and similar in the Std-O_2-_, Thick-O_2+_, and Thick-O_2-_ conditions compared to the Std-O_2+_, Std-P2O, and Thick-P2O conditions, we concluded that the former set of conditions were anoxic. The Thick-O_2+_ and the Std-O_2-_ devices had almost identical relative changes in quantified growth, meaning that the Thick-O_2+_ device under normal oxygen (normoxic) successfully maintained the same condition as the Std-O_2-_ condition (2.5mm top cover under anoxic condition).

The fact that Std-O_2-_, Thick-O_2+_ and Thick-O_2-_ conditions established a hypoxic environment was further supported by testing the growth of *E. coli* Nissle 1917 (ECN) (**Figure 4C** and **4D**). ECN is a facultative anaerobe, which means that its growth will be higher under normoxic rather than under anoxic conditions. ECN growth was supported only by the Std-O_2+_ condition (**Figure 4D**); there were far fewer bacteria in the Std-O_2-_, Std-P2O, Thick-O_2+_, Thick-O_2-_, and Thick-P2O conditions compared to the Std-O_2+_ condition. Further, the growth of ECN is very similar to the growth in Std-O_2-_, Thick-O_2+_, and Thick-O_2-_ conditions (**Figure 4D, Table S2**) and significantly lowerin Std-O_2+_; only the Std-O_2+_ condition supported the bacteria’s growth (**Figures 4E** and **4F, Table S3**), suggesting that the Std-O_2+_ condition modeled the normoxic condition. Since there was a lack of growth in other devices, namely in the thick device, it can be concluded that anoxic environment was created in these devices. In addition, since the difference of relative increases in bacteria in (**Figure 4D**) and mean fluorescence intensity (**Figure 4F**) between the Thick-O_2+_ and Std-O_2-_ conditions were not statistically significant while being significantly different from the Std-O_2+_, we can conclude that an anoxic environment was established in the Thick-O_2+_ condition (5mm top cover in environment with normal oxygen).

Surprisingly, all the tested bacteria in the device with P2O conditioned media in top reservoir showed lower growth (*B. adolescentis* (**Figure 4A and 4B**), *E. Coli* Nissle 1917 (**Figure 4C and 4D**), and GFP-ECN (**Figure 4E and 4F**)). GFP-ECN congregated together instead of growing into a biofilm, suggesting that the oxygen scavenger interfered with bacteria growth. To conclude, the above results validate that the device with a thick PDMS top cover (5mm) supports the formation of anoxic environment in the top chamber even when placed in an environment with normal oxygen.

To confirm the oxygenation status of the cells, we tested all the devices to see whether the cells were hypoxic when the oxygen environment was established. We treated the cells with pimonidazole which, in hypoxic conditions, is reduced to form stable covalent adducts with thiol groups that are fluorescent. We found that from the start of the experiment to around 4 hours, cells in all of the devices had similar oxygenation (**Figure 5A, Figure S5**). At 8hrs, the cells in Thick-O_2+_, Thick-O_2-_, and the Std-O_2-_ conditions had more visible fluorescence (indicating hypoxia) compared to those in the Std-O_2+_ condition (**Figure 5A**, **Figure S5**). By 24h, hypoxia was visible in cells in the Thick-O_2+_, Thick-O_2-_, and the Std-O_2-_ conditions compared to the Std-O_2+_. The normalized fluorescence intensity per nuclei in the Thick-O_2+_, Thick-O_2-_, and Std-O_2-_ conditions was almost double that for the cells in the Std-O_2+_ condition (**Figure 5B**, **Table S4**). These results indicate that over the progression of 24 hours, epithelial cells in the Std-O_2-_, Thick-O_2+_, Thick-O_2-_ had very similar intensity with no statistical difference, however, those in the Thick-O_2+_, Thick-O_2-_, and Std-O_2-_ conditions showed a significant higher intensity compared to Std-O_2+_ suggesting the presence of an anoxic environment.

### 3.3 Caco-2 cells functional changes in hypoxic devices

ZO-1 antibody staining was used to show the tight junctions representing the integrity of the epithelial cell monolayer. The tight junctions are lined between the epithelial cells (Dörfel and Huber, 2012) and necessary for these intestinal cells to properly function (Lee, 2015; Goyer et al., 2016; Chelakkot et al., 2018).

In Std-O_2+_ condition, which served as the control, over the course of 24-hours, there were no changes in the ZO-1 protein (**Figure 6A**). In contrast, ZO-1 was downregulated in cells in the Thick-O_2-_, Std-O_2-_and Thick-O_2+_ conditions over the 24-hour period (**Figure 6B – 6D**), confirming that the lack of oxygen disrupted the tight junctions (**Figure 6E**, **Table S5**). Our results are supported by other research which shows that the hypoxia of cells increased monolayer permeability, and lowered TEER values (Page et al., 2016; Xu et al., 1999). This data further supports that a device with thicker cover (Thick-O_2+_) establishes a hypoxic environment.

**Figure 6:**
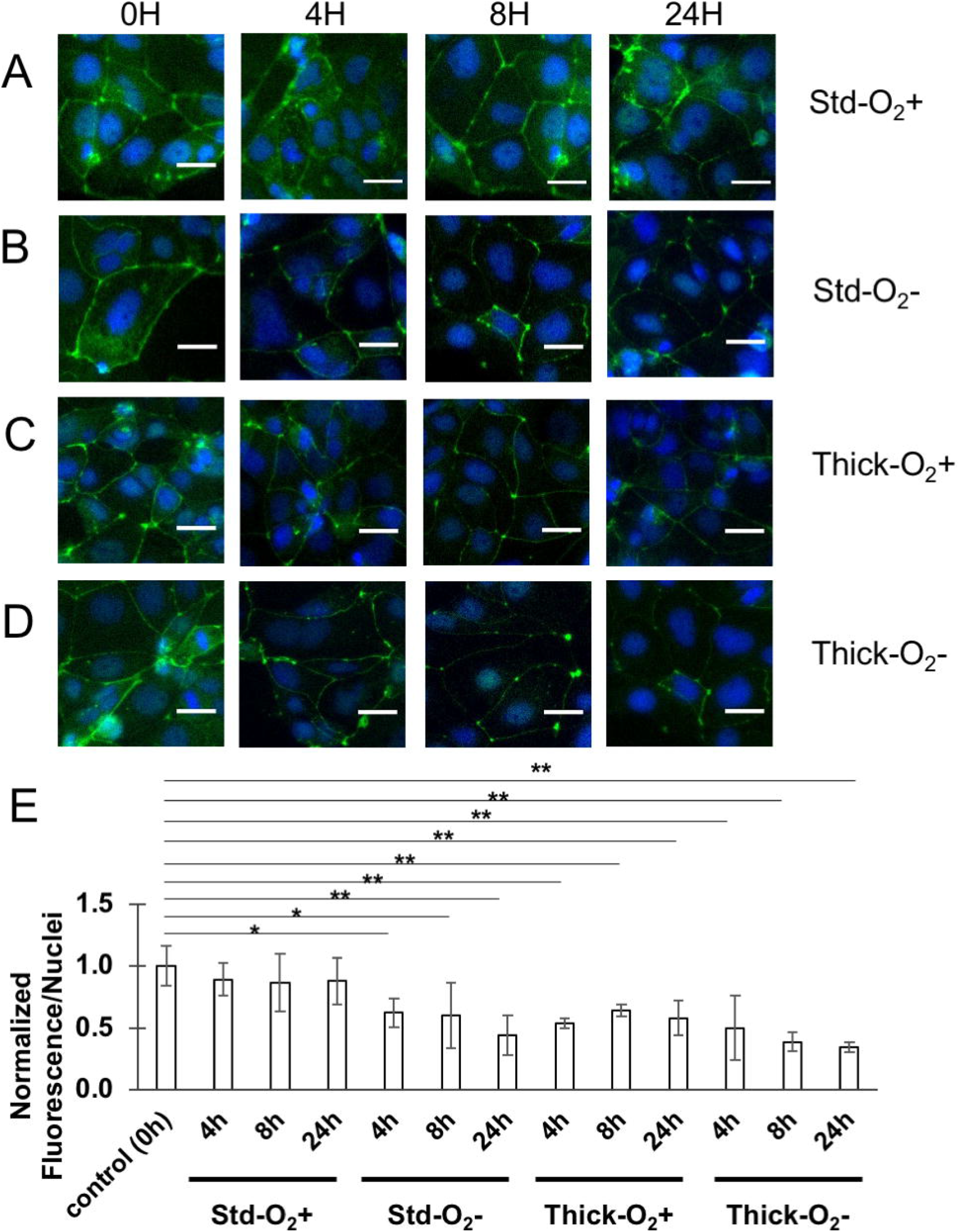
ZO-1 immunostaining in Std-O_2+_ (A), Std-O_2-_ (B), Thick-O_2+_ (C), and Thick-O_2-_ (D) when incubated for 0-24 hours. Scale bar is 20 μm. (E) Quantification of changes in tight junction protein ZO-1 in the different conditions. *: *p*-value < 0.05; **: *p*-value < 0.01.

### 3.4 Control of oxygen levels in 2-chamber device

As discussed before, we used three different perfusion flow rates (5 μL/h, 7.2 μL/h and 30 μL/h) to test the oxygenation of the cells as well as simultaneous formation of anoxia in the top chamber. We incorporated oxygen sensors in the top chamber to monitor oxygen changes during bacteria and Caco-2 cells coculture (**Figure 7A-C**). The fluid in the top chamber was static whereas the fluid in the bottom chamber was perfused. After 24h of culture, the oxygen concentration of top chamber in each condition was calculated (**Table 4**) against a calibration curve (**Figure S3**). We found that at 5 μL/h and 7.2 μL/h, the sensors recorded oxygen levels at 0.03 ± 0.07% and 0.08 ± 0.013% respectively. In contrast, at 30 μL/h, the oxygen levels increased to 12.98 ± 5.85%. These results indicate that when at a low flow rate (5 and 7.2 μL/h), the top chamber was anoxic; but when the flow rate is high (30 μL/h), the top chamber was normoxic.

**Figure 7:**
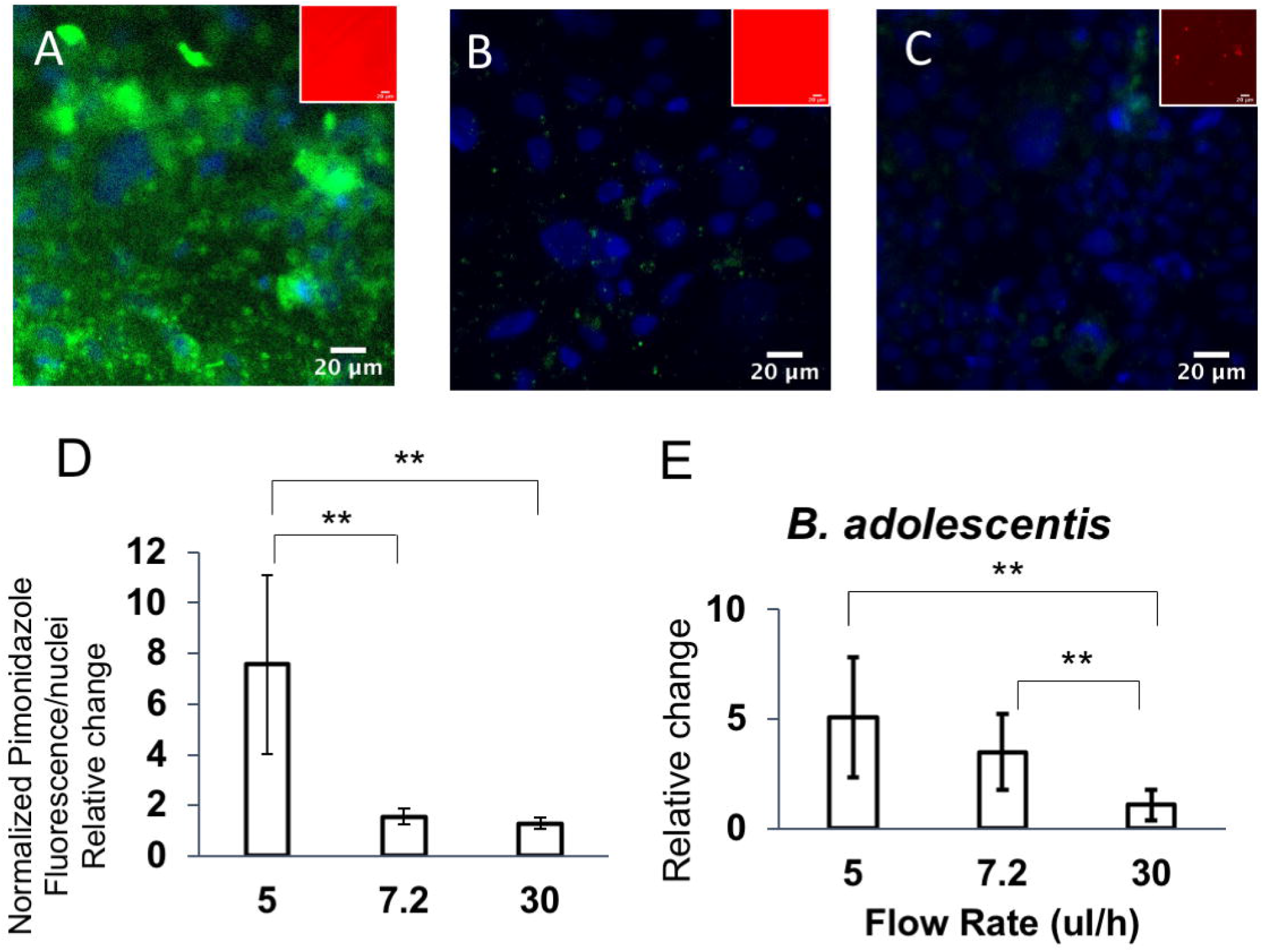
(A-C) Immunostained images of Caco-2 cells cocultured with *B. adolescentis* in thick 2-chamber device with different flow rates – 5 μL/h (A), 7.2 μL/h (B) and 30 μL/h (C). Scale bar is 20 μm. Inset shows the fluorescence of the oxygen sensor. (D) Quantification of pimonidazole fluorescence. (E) Relative growth rate of *B. adolescentis* in the 2-chamber device. **: p value <0.01; *: p value <0.05; n.s.: no significance.

**Table 4.** Oxygen concentration in top chamber of the 2-chamber device as measured by the oxygen sensor.

| **Groups** | **O<sub>2</sub> Concentration (%)** | **SEM (n≥18)** |
| --- | --- | --- |
| **5 ul/h** | 0.03 | 0.07 |
| **7.2 ul/h** | 0.07 | 0.14 |
| **30 ul/h** | 13.52 | 6.1 |

We confirmed these findings by testing the hypoxic/normoxia of Caco-2 cells and the growth of anaerobic bacteria, *B. adolescentis* (**Figure 7**). The hypoxia of Caco-2 cells was quantified by measuring the fluorescence (**Figure 7D**) and the growth of *B. adolescentis* was measured and shown in **Figure 7E**. The results show that the cells are hypoxic in the device with a flow rate of 5 μL/h (**Figure 7A** and **7D**). At 7.2 μL/h, Caco-2 cells did not show hypoxia (**Figure 7B**) and supported growth of *B. adolescentis* (**Figure 7E**). At 30 μL/h, Caco-2 cells did not show hypoxia as well (**Figure 7C**) and as expected based on oxygen sensor (**Table 4**), the device provided less ideal conditions for *B. adolescentis’* growth.

To determine the growth of anaerobes, *B. adolescentis* was cocultured with Caco-2 monolayers in the same top chamber. After 24h of coculture, the bacteria were eluted out, collected, and cultured on an agar plate. The initial seeding density of *B. adolescentis* in devices was kept the same. The growth of bacteria in each device was quantified and the relative changes of bacteria growth in device was normalized according to the initial seeding density. We found that the growth of *B. adolescentis* in the device with 30 μL/h flow rate was much lower than the growth in the devices with flow rates of 5 μL/h and 7.2 μL/h. We surmised that more oxygen was present within the device under higher flow rates, which inhibited the bacteria’s growth; in other words, the top chamber was hypoxic at lower flow rates.

Based upon these flow rates, we estimated the oxygen consumption rate of the Caco-2 cells when cultured in the devices (**Table 5**). Interestingly, our values were much lower than those reported in the literatures. Some recent studies have indicated that in vitro studies may have overestimated the oxygen consumption rates that are observed in vivo (Ahluwalia, 2017),(Tirella et al., 2015). This has to do with the oxygen supply in vivo being limited by the oxygen transport from the capillaries to the cells (which we have mimicked), rather than being always available as is modeled in traditional tissue culture setups. Our experiment can serve as a model to study oxygen consumption of cells under different physiological conditions, such as under interactions by different bacterial species.

**Table 5.** Oxygen consumption rate for the different perfusion flow rate.

| Perfusion flow rate $Q$ ($\mu\text{L/h}$) | Oxygen consumption rate $q$ ($\text{mol/s} \cdot \text{cell}$) of the cells |
| --- | --- |
| 5 | $1.35 \times 10^{-19}$ |
| 7.2 | $1.95 \times 10^{-19}$ |
| 30 | $8.16 \times 10^{-19}$ |

To summarize, we have experimentally established the relationship between media flow rate with the oxygen concentration in the top chamber where bacteria and intestinal epithelial cells are cocultured. We found that in our 2-chamber device, a flow rate of 7.2 μL/h supported oxygenation of Caco-2 cells while simultaneously providing an anoxic environment for bacteria growth in the top chamber. The results indicate that at flow rates lower than this, the cells become hypoxic. Conversely, at higher flow rates, the cells are oxygenated but the top chamber becomes normoxic. Our experimental model is supported by the mathematical model that considers diffusion of oxygen into the top chamber and the cellular oxygen consumption rate.

## 4. Conclusions

The intestinal microenvironment consists of the epithelium along with the microbiota under different oxygen tensions. The oxygen concentration in the small intestine and colon ranges from 1 to 30 mmHg, respectively (Sheridan et al., 1990). Due to the unique oxygenation profile in the intestine, there is a strong need for developing representative and simple experimental models. In this paper, we created standalone microfluidic devices that formed dual-oxygen environment without the use of an external anaerobic chamber or oxygen scavengers. Our work established a simple and precise method to control oxygen levels to coculture intestinal epithelial and bacterial cells. We showed that by changing the thickness of the device cover, the oxygen tension in the chamber could be modulated. In addition, we found that the oxygen tension in the cell chamber could be controlled by controlling flow rate in the bottom chamber. Our experimental model is supported by the mathematical model that considers diffusion of oxygen into the top chamber and the cellular oxygen consumption rate.

Previous approaches have either required placing the microfluidic organ-chip device into an anaerobic chamber or required continuous perfusion of the bacteria. Our work is a significant improvement over existing approaches because the devices are standalone and provide a more accurate representation of the intestinal environment. This enabled us to calculate oxygen consumption rate of the cells. We verified the oxygen levels using a number of tests: microscale oxygen sensitive sensors incorporated within the devices, hypoxic immunostaining of Caco-2 cells, and genetically encoded bacteria. Collectively, these methods monitored oxygen concentrations in device more comprehensively than previous reports and allowed for control of oxygen tension to match the requirements of both intestinal cells and anaerobic bacteria. This method we developed can be broadly applied to mimicking multiple physiological scenarios where oxygen tension varies including the anaerobic/aerobic microbiota-cells coculture, the cancer hypoxia microenvironment, and luminal microenvironment of the intestine. Furthermore, this method to control oxygen tension can be utilized to control other chemical elements when a heterogenous environment is required within same system, such as modulation of glucose and growth factors.

## Acknowledgements

This work was supported in part by the Alternatives Research and Development Foundation, Starr and Fieldhouse Fellowship, and student scholarships from the Armor College of Engineering, Illinois Institute of Technology. We acknowledge technical help from Dr. Seok Hoon Hong in constructing the *E.coli* Nissle 1917 with GFP.

## Notes

### Competing Interest Statement

The authors have declared no competing interest.

