## Supplementary material for "A Novel Standalone Microfluidic Device for Local Control of Oxygen Tension for Intestinal-Bacteria Interactions": Figure S1-S5

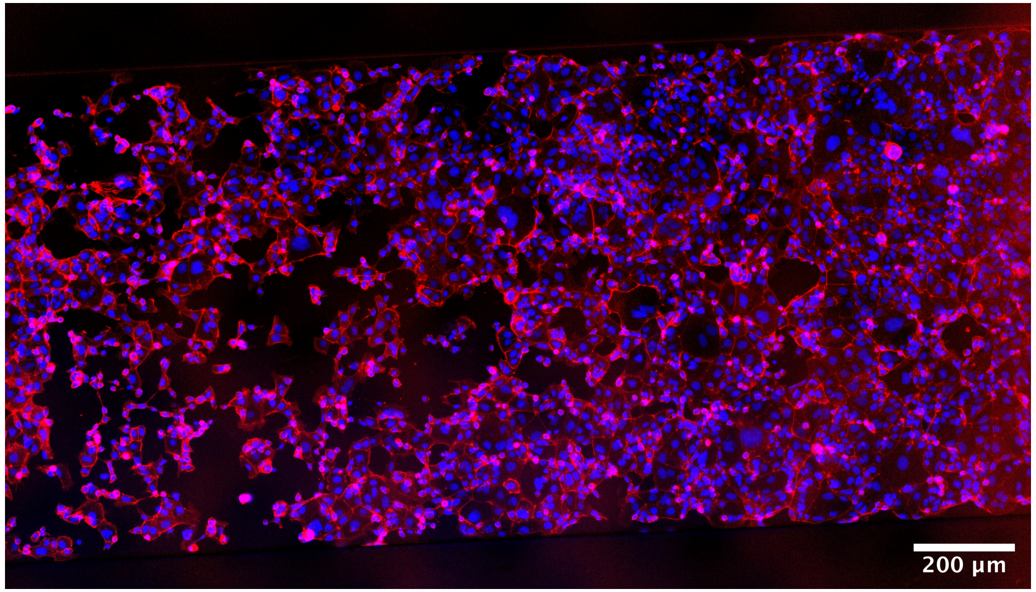


**Figure S1.** F-actin (Red) and Hoechst (blue) stained Caco-2 cells. Cells were cultured in microfluidic device for 4 days. Images were taken at day 4 and to show a fully confluent monolayer of intestinal epithelial cells covered along the device. Scale bar: 200 µm.

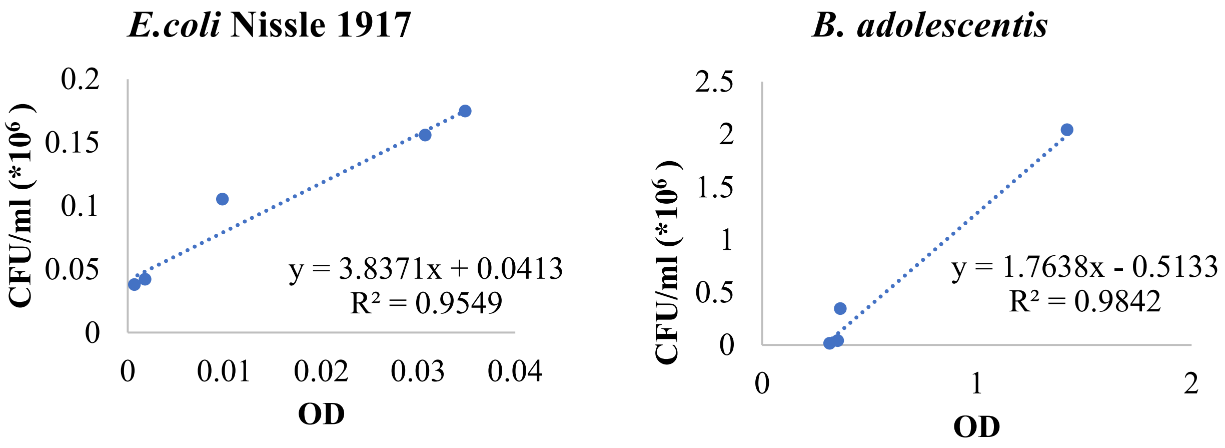


**Figure S2**. Bacteria standard curve of OD and density (CFU/mL). n=6.

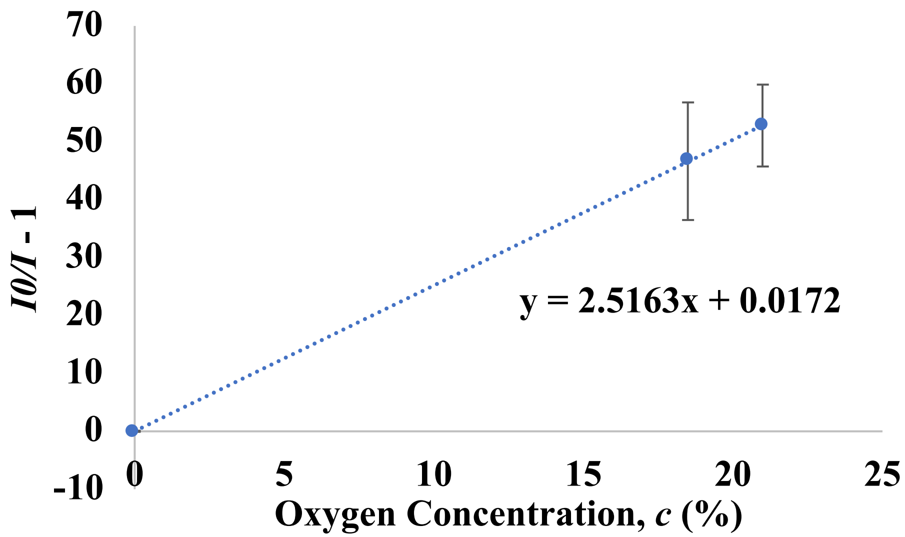


**Figure S3**. $O_2$ sensor calibration, n>= 6.

A

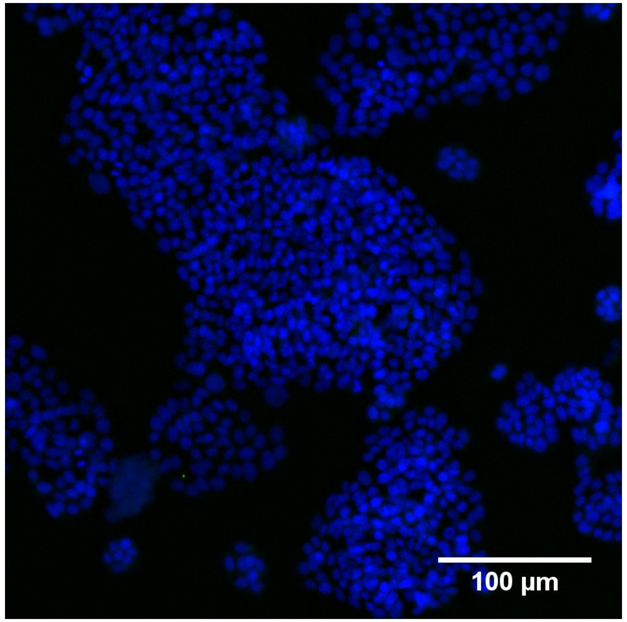


B

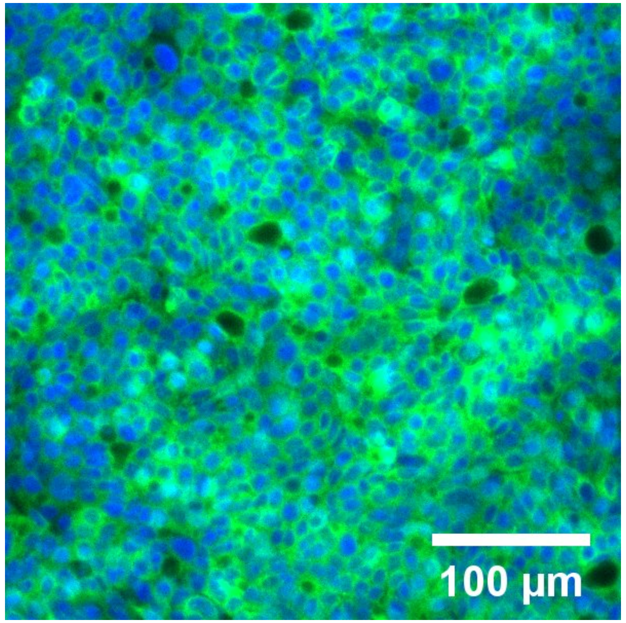


**Figure S4. Validation of cellular hypoxia in tissue culture plate with different oxygen tension culture environment.** Caco-2 cells were cultured in tissue culture plate in standard tissue culture incibator till 80% confluency and then put the plates in different environment: standard tissue culture incubator (A) and anaerobic jar (B). After 24 hours, cells were treated with 200 µM pimonidazole HCl (Hypoxyprobe) for 2 hours to determine hypoxia. Then the cells were fixed and stained with FITC-Mab (Hypoxyprobe) and FITC HRP MAb (Hypoxyprobe) and Hoechst (ThermoFisher). The result of staining was then imaged using fluorescent microscope.

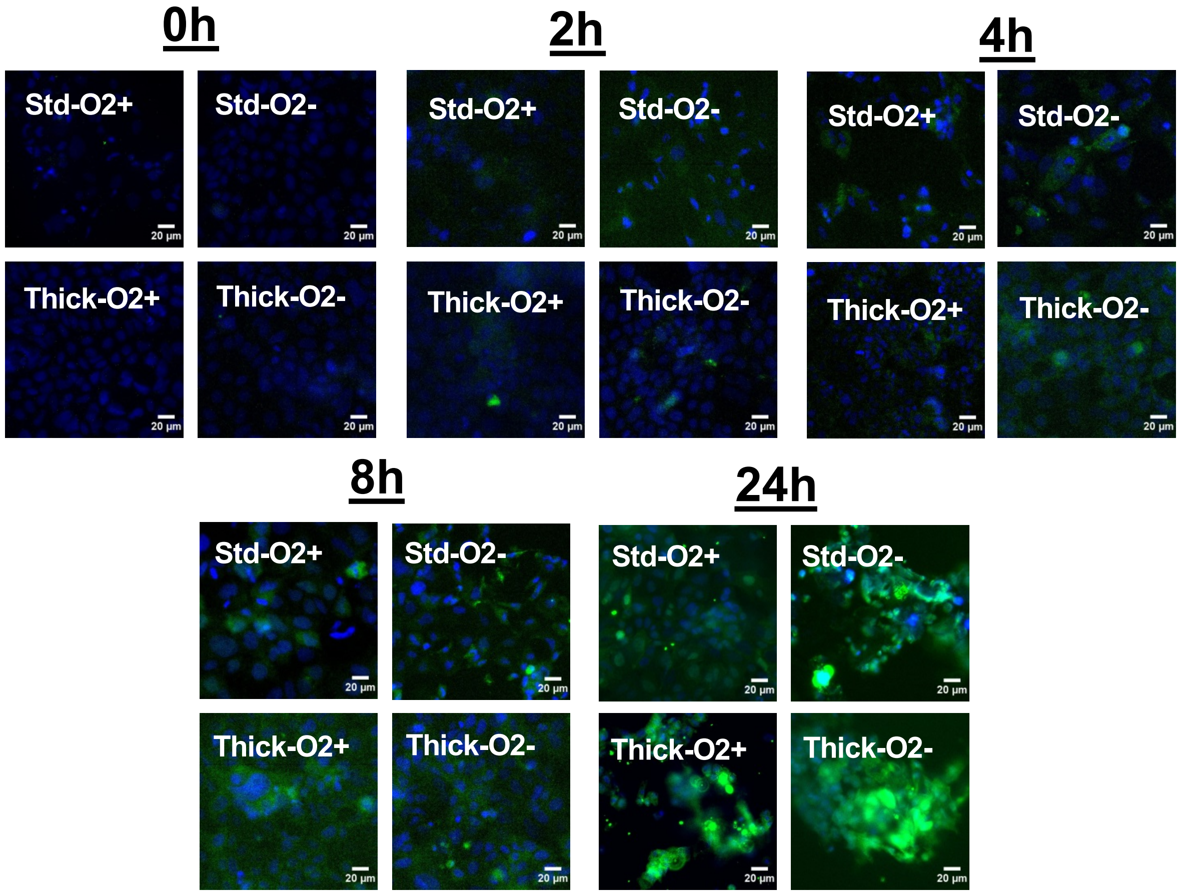


**Figure S5.** Hypoxia immunostained images of normoxic and hypoxic Caco-2 cells in devices with standard and thick top cover for different treatment intervals: 0, 2, 4, 8 and 24 hours, using pimonidazole staining kit. Green: hypoxia; Blue: nuclei. Scale bar is 20 µm.
