## Supplementary material for "A Novel Standalone Microfluidic Device for Local Control of Oxygen Tension for Intestinal-Bacteria Interactions": Table S1-S6

| **NORMALIZED DATA** | **Std-$O_2^+$** | **Std-$O_2^-$** | **Std-P2O** | **Thick-$O_2^+$** | **Thick-$O_2^-$** | **Thick-P2O** |
|---|---|---|---|---|---|---|
| **Mean** | 1.134 | 18.346 | 1.613 | 19.285 | 19.819 | 1.898 |
| **SEM (n≥6)** | 0.600 | 6.693 | 0.809 | 4.331 | 6.280 | 1.280 |

**p-value (t-test)**

| | | | | | | | | | |
|---|---|---|---|---|---|---|---|---|---|
| Std-$O_2^+$ vs Std-$O_2^-$** | 0.000 | Std-$O_2^-$ vs Std-P2O** | 0.000 | **Std-P2O** vs **Thick-$O_2^+$**** | 0.000 | Thick-$O_2^+$ vs Thick-$O_2^-$ | 0.801 | Thick-$O_2^-$ vs Thick-P2O** | 0.000 |
| Std-$O_2^+$ vs Std-P2O | 0.127 | **Std-$O_2^-$** vs **Thick-$O_2^+$** | 0.657 | Std-P2O vs Thick-$O_2^-$** | 0.000 | Thick-$O_2^+$ vs Thick-P2O** | 0.000 | | |
| **Std-$O_2^+$** vs **Thick-$O_2^+$**** | 0.000 | Std-$O_2^-$ vs Thick-$O_2^-$ | 0.539 | Std-P2O vs Thick-P2O | 0.519 | | | | |
| Std-$O_2^+$ vs Thick-$O_2^-$** | 0.000 | Std-$O_2^-$ vs Thick-P2O** | 0.000 | | | | | | |
| Std-$O_2^+$ vs Thick-P2O | 0.087 | | | | | | | | |

**Table S1.** *B. adolescentis* growth in 1-chamber device with standard or thick PDMS top cover, in normoxic or hypoxic incubation conditions; and statistical analysis between different experimental conditions. * p-value < 0.05/6 =0.008 (Bonferroni correction for n multiple significance tests) ; ** p-value < 0.01/6 = 0.0017.

| NORMALIZED DATA | Std-$O_2^+$ | Std-$O_2^-$ | Std-P2O | Thick-$O_2^+$ | Thick-$O_2^-$ | Thick-P2O |
|---|---|---|---|---|---|---|
| **Mean** | 40.675 | 5.430 | 6.051 | 4.220 | 4.098 | 4.649 |
| **SEM (n≥6)** | 12.541 | 1.770 | 2.364 | 1.088 | 1.611 | 1.615 |

**p-value (t-test)**

| | | | | | | | | | |
|---|---|---|---|---|---|---|---|---|---|
| Std-$O_2^+$ vs Std-$O_2^-$** | 0.000 | Std-$O_2^-$ vs Std-P2O | 0.463 | **Std-P2O** vs **Thick-$O_2^+$** | 0.027 | Thick-$O_2^+$ vs Thick-$O_2^-$ | 0.831 | Thick-$O_2^-$ vs Thick-P2O | 0.412 |
| Std-$O_2^+$ vs Std-P2O** | 0.000 | **Std-$O_2^-$** vs **Thick-$O_2^+$** | 0.045 | Std-P2O vs Thick-$O_2^-$ | 0.029 | Thick-$O_2^+$ vs Thick-P2O | 0.455 | | |
| **Std-$O_2^+$** vs **Thick-$O_2^+$*** ** | 0.000 | Std-$O_2^-$ vs Thick-$O_2^-$ | 0.056 | Std-P2O vs Thick-P2O | 0.106 | | | | |
| Std-$O_2^+$ vs Thick-$O_2^-$** | 0.000 | Std-$O_2^-$ vs Thick-P2O | 0.251 | | | | | | |
| Std-$O_2^+$ vs Thick-P2O** | 0.000 | | | | | | | | |

**Table S2**. ECN Bacteria growth in 1-chamber device with standard or thick PDMS top cover, in normoxic or hypoxic incubation conditions; and statistical analysis between different experimental conditions. * p-value < 0.05/6 =0.008 (Bonferroni correction for n multiple significance tests) ; ** p-value < 0.01/6 = 0.0017.

| NORMALIZED DATA | Std-$O_2^+$ | Std-$O_2^-$ | Std-P2O | Thick-$O_2^+$ | Thick-$O_2^-$ | Thick-P2O |
|---|---|---|---|---|---|---|
| **Mean** | 62.08 | 10.573 | 11.815 | 12.489 | 12.497 | 13.698 |
| **SEM (n≥6)** | 8.964 | 2.109 | 2.170 | 2.145 | 1.222 | 2.519 |

| p-value (t-test) | | | | | | | | | |
|---|---|---|---|---|---|---|---|---|---|
| Std-$O_2^+$ vs Std-$O_2^-$** | 0.000 | Std-$O_2^-$ vs Std-P2O | 0.284 | **Std-P2O** vs **Thick-$O_2^+$** | 0.520 | Thick-$O_2^+$ vs Thick-$O_2^-$ | 0.424 | Thick-$O_2^-$ vs Thick-P2O | 0.181 |
| Std-$O_2^+$ vs Std-P2O** | 0.000 | **Std-$O_2^-$** vs **Thick-$O_2^+$** | 0.123 | Std-P2O vs Thick-$O_2^-$ | 0.705 | Thick-$O_2^+$ vs Thick-P2O | 0.401 | | |
| **Std-$O_2^+$** vs **Thick-$O_2^+$**** | 0.000 | Std-$O_2^-$ vs Thick-$O_2^-$ | 0.674 | Std-P2O vs Thick-P2O | 0.196 | | | | |
| Std-$O_2^+$ vs Thick-$O_2^-$** | 0.000 | Std-$O_2^-$ vs Thick-P2O | 0.059 | | | | | | |
| Std-$O_2^+$ vs Thick-P2O** | 0.000 | | | | | | | | |

**Table S3.** ECN-GFP Bacteria growth in 1-chamber device (confocal microscope) with standard or thick PDMS top cover, in normoxic or hypoxic incubation conditions; and statistical analysis between different experimental conditions. * p-value < 0.05/6 =0.008 (Bonferroni correction for n multiple significance tests) ; ** p-value < 0.01/6 = 0.0017.

| Time intervals | 2h | | | | 4h | | | | 8h | | | | 24h | | | |
|---|---|---|---|---|---|---|---|---|---|---|---|---|---|---|---|---|
| Groups | Std-$O_2^+$ | Std-$O_2^-$ | Std-P2O | Thick-$O_2^+$ | Std-$O_2^+$ | Std-$O_2^-$ | Std-P2O | Thick-$O_2^+$ | Std-$O_2^+$ | Std-$O_2^-$ | Std-P2O | Thick-$O_2^+$ | Std-$O_2^+$ | Std-$O_2^-$ | Std-P2O | Thick-$O_2^+$ |
| **Mean** | 0.583 | 1.101 | 0.575 | 0.555 | 1.641 | 1.930 | 1.512 | 1.824 | 2.128 | 2.994 | 3.172 | 3.483 | 3.423 | 6.422 | 6.565 | 6.194 |
| **SEM (n≥6)** | 0.334 | 0.127 | 0.110 | 0.345 | 0.219 | 0.436 | 0.360 | 0.528 | 0.389 | 0.934 | 0.726 | 1.027 | 0.696 | 0.962 | 0.916 | 1.643 |

**2h p-value (t-test)**

| | | | | | |
|---|---|---|---|---|---|
| Std-$O_2^+$ vs Std-$O_2^-$ | 0.023 | Std-$O_2^-$ vs Thick-$O_2^+$ * | 0.006 | Thick-$O_2^+$ vs Thick-$O_2^-$ | 0.884 |
| Std-$O_2^+$ vs Thick-$O_2^+$ | 0.962 | Std-$O_2^-$ vs Thick-$O_2^-$ * | 0.004 | | |
| Std-$O_2^+$ vs Thick-$O_2^-$ | 0.885 | | | | |

**4h p-value (t-test)**

| | | | | | |
|---|---|---|---|---|---|
| Std-$O_2^+$ vs Std-$O_2^-$ | 0.383 | Std-$O_2^-$ vs Thick-$O_2^+$ | 0.273 | Thick-$O_2^+$ vs Thick-$O_2^-$ | 0.452 |
| Std-$O_2^+$ vs Thick-$O_2^+$ | 0.630 | Std-$O_2^-$ vs Thick-$O_2^-$ | 0.804 | | |
| Std-$O_2^+$ vs Thick-$O_2^-$ | 0.623 | | | | |

**8h p-value (t-test)**

| | | | | | |
|---|---|---|---|---|---|
| Std-$O_2^+$ vs Std-$O_2^-$ | 0.245 | Std-$O_2^-$ vs Thick-$O_2^+$ | 0.808 | Thick-$O_2^+$ vs Thick-$O_2^-$ | 0.693 |
| Std-$O_2^+$ vs Thick-$O_2^+$ | 0.114 | Std-$O_2^-$ vs Thick-$O_2^-$ | 0.575 | | |
| Std-$O_2^+$ vs Thick-$O_2^-$ | 0.137 | | | | |

**24h p-value (t-test)**

| | | | | | |
|---|---|---|---|---|---|
| Std-$O_2^+$ vs Std-$O_2^-$ * | 0.003 | Std-$O_2^-$ vs Thick-$O_2^+$ | 0.827 | Thick-$O_2^+$ vs Thick-$O_2^-$ | 0.710 |
| Std-$O_2^+$ vs Thick-$O_2^+$ * | 0.004 | Std-$O_2^-$ vs Thick-$O_2^-$ | 0.816 | | |
| Std-$O_2^+$ vs Thick-$O_2^-$ * | 0.010 | | | | |

**Table S4.** Caco-2 cells hypoxia test in 1-chamber device with standard or thick PDMS top cover, in normoxic or hypoxic incubation conditions, in different time point; and statistical analysis between different experimental conditions. * p-value < 0.05/4 =0.0125 (Bonferroni correction for n multiple significance tests) ; ** p-value < 0.01/4 = 0.0025.

| Groups | Control | Std-$O_2^+$ | | | Std-$O_2^-$ | | | Thick-$O_2^+$ | | | Thick-$O_2^-$ | | |
|---|---|---|---|---|---|---|---|---|---|---|---|---|---|
| Time intervals | 0h | 4h | 8h | 24h | 4h | 8h | 24h | 4h | 8h | 24h | 4h | 8h | 24h |
| Mean | 1.000 | 0.891 | 0.864 | 0.877 | 0.622 | 0.599 | 0.442 | 0.538 | 0.641 | 0.579 | 0.499 | 0.388 | 0.343 |
| SEM (n≥6) | 0.162 | 0.133 | 0.230 | 0.185 | 0.114 | 0.263 | 0.159 | 0.039 | 0.051 | 0.141 | 0.261 | 0.074 | 0.042 |

p-value (t-test)

| Std-$O_2^+$ | | | | | | Std-$O_2^-$ | | | | | |
|---|---|---|---|---|---|---|---|---|---|---|---|
| 0h vs 4h | 0.484 | 4h vs 8h | 0.846 | 8h vs 24h | 0.977 | 0h vs 4h* | 0.004 | 4h vs 8h | 0.553 | 8h vs 24h | 0.149 |
| 0h vs 8h | 0.254 | 4h vs 24h | 0.798 | | | 0h vs 8h* | 0.005 | 4h vs 24h* | 0.003 | | |
| 0h vs 24h | 0.113 | | | | | 0h vs 24h** | 0.000 | | | | |
| **Thick-$O_2^+$** | | | | | | **Thick-$O_2^-$** | | | | | |
| 0h vs 4h** | 0.000 | 4h vs 8h | 0.017 | 8h vs 24h | 0.134 | 0h vs 4h** | 0.001 | 4h vs 8h | 0.323 | 8h vs 24h | 0.180 |
| 0h vs 8h** | 0.000 | 4h vs 24h | 0.857 | | | 0h vs 8h** | 0.000 | 4h vs 24h | 0.257 | | |
| 0h vs 24h** | 0.000 | | | | | 0h vs 24h** | 0.000 | | | | |

**Table S5.** Caco-2 function changes in device with standard or thick PDMS top cover, in normoxic or hypoxic incubation conditions, in different time point; and statistical analysis between different experimental conditions. * p-value < 0.05/4 =0.0125 (Bonferroni correction for n multiple significance tests) ; ** p-value < 0.01/4 = 0.0025.

| Flow rate (µL/h ) | 5 | 7.2 | 30 |
| --- | --- | --- | --- |
| **Mean** | 6.56 | 0.56 | 0.29 |
| **SEM (n≥6)** | 3.55 | 0.31 | 0.22 |

**p-value (t-test)**

| | | | |
| --- | --- | --- | --- |
| **5 ul/h-7.2 ul/h ** ** | 0.0032 | **7.2 ul/h-30 ul/h** | 0.0734 |
| **5 ul/h-30 ul/h ** ** | 0.0026 | | |

**Table S7.** Caco-2 cells hypoxia (pimonidazole) test in 2-chamber coculture device with thick PDMS top cover, in normoxic incubation conditions, with different bottom media flow rate; and statistical analysis between different flow rate. * p-value < 0.05/3 =0.017 (Bonferroni correction for n multiple significance tests) ; ** p-value < 0.01/3 = 0.0033.

| Groups | 5 ul/h | 7.2 ul/h | 30 ul/h |
|---|---|---|---|
| **Mean** | 4.93 | 3.49 | 1.09 |
| **SEM (n≥6)** | 1.69 | 0.81 | 0.33 |

| p-value (t-test) | | | |
|---|---|---|---|
| **5 ul/h-7.2 ul/h** | 0.144 | **7.2 ul/h-30 ul/h **** | 0.003 |
| **5 ul/h-30 ul/h **** | 0.001 | | |

**Table S6.** *B. adolescentis* growth in 2-chamber coculture device with thick PDMS top cover, in normoxic incubation conditions, with different bottom media flow rate; and statistical analysis between different flow rate. * p-value < 0.05/3 =0.017 (Bonferroni correction for n multiple significance tests) ; ** p-value < 0.01/3 = 0.0033.
